# Urothelial organoids originate from Cd49f^High^ stem cells and display Notch-dependent differentiation capacity

**DOI:** 10.1101/287979

**Authors:** Catarina P. Santos, Eleonora Lapi, Laura Álvaro-Espinosa, Asunción Fernández-Barral, Antonio Barbáchano, Diego Megías, Alberto Muñoz, Francisco X Real

## Abstract

The urothelium is a specialized stratified epithelium with unique structural and functional features. Understanding the mechanisms involved in urothelial stem cell biology and differentiation has been limited by the lack of methods for unlimited propagation. Here, we establish normal mouse urothelial organoid (NMU-o) cultures that can be maintained uninterruptedly for >1 year. Organoid growth is dependent on EGF and Wnt activators. High CD49f/ITGA6 expression features a subpopulation of organoid-forming urothelial stem cells expressing basal markers. On induction of differentiation, multilayered organoids show reduced layer number, acquire barrier function, and activate the urothelial program, including expression of uroplakins and tight junction components. Combined pharmacological modulation of PPARγ and EGFR was most potent driving cell differentiation. Transcriptome analysis of organoids widely validates the system, highlights the transcriptional networks involved, and reveals NOTCH signaling as a novel pathway required for normal urothelial organoid differentiation.

## Introduction

Urinary bladder diseases, most notably cystitis and bladder cancer, are important medical problems that generate high costs to the health systems worldwide. The bladder, ureters, renal pelvis, and part of the urethra are lined by the urothelium, a stratified epithelium reminiscent of the skin epidermis consisting of three cell types organized in 3-7 layers (Figure 1A). Basal cells are small, cuboid, and express CD44 and KRT5; a fraction thereof express KRT14 and have stem cell properties (Papafotiou et al., 2016). Intermediate cells are larger, express KRT8, KRT18, and uroplakins UPK1a, 1b, 2, 3a and 3b. Luminal umbrella cells are largest, multinucleated, highly specialized cells expressing high levels of uroplakins and KRT20 (Hicks, 1965; Ho et al., 2012; Wu et al., 1990). Umbrella cells constitute the physiological barrier to the passage of water, electrolytes, and urea through tight junctions, responsible for the high resistance paracellular pathway (Apodaca, 2004). Unlike the skin epidermis, the urothelium has a very slow turnover (Jost, 1986) yet it preserves a robust capacity to restore epithelial integrity upon damage (Claysin and Cooper, 1969; Papafotiou et al., 2016). Several key transcriptional factors involved in urothelial proliferation/differentiation have been identified using 2D cultures and genetic mouse models. PPARγ is expressed in the urothelium throughout embryonic development and in the adult (Jain et al., 1998) and it has been shown to participate in proliferation, cooperating with FOXA1, KLF5, and EGFR signaling (Bell et al., 2011; Varley et al., 2009). The role of these pathways in urothelial differentiation is incompletely understood, due to the lack of methods to continually propagate normal cells.

**Figure 1.**
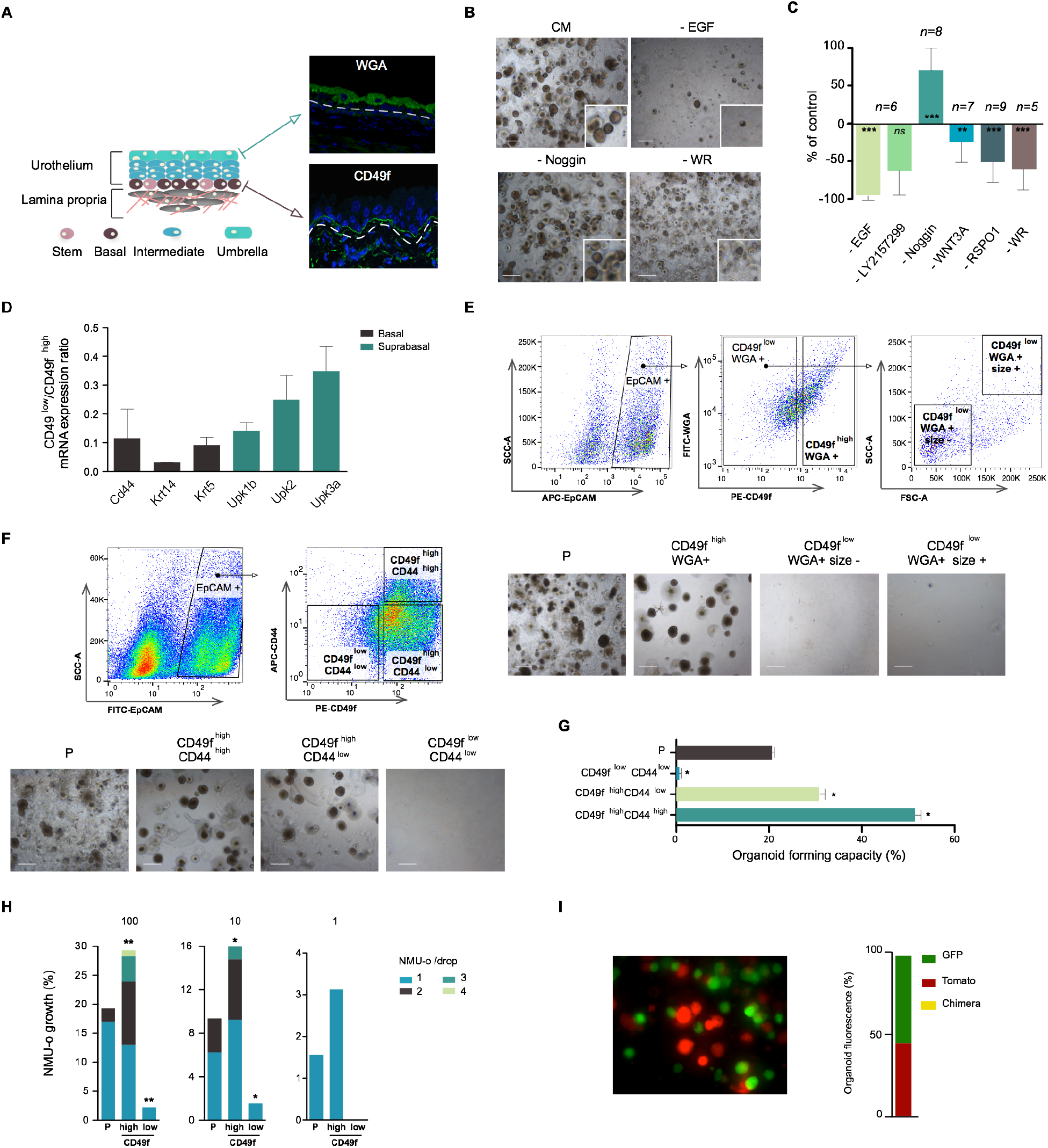
CD49f labels an organoid-forming mouse urothelial cell population with stem cell features. (**A**) Urothelium cyto-organization highlighting the basal (CD49f) and luminal (WGA-binding) markers. (**B**) Representative images of organoids from urothelial cell suspensions embedded in Matrigel in CM (upper left). The remaining panels correspond to the “Leave-one-out” experiments (see C). (**C**) Quantification of NMU-o growth in “Leave-one-out” experiments (WR condition: WNT3A and RSPO1 were omitted). Number of organoids referred to CM; error bars indicate SEM. (**D**) Enrichment of urothelial marker transcripts in CD49f^low^ cells. Graph representative of one experiment out of 2 and error bars indicate SD. (**E, F**) FACS plots depicting the analysis and isolation of EpCAM^+^ cells according to cell surface markers and size (n=2). (**G**) Quantification of the organoid-forming capacity of sorted urothelial cells, compared to unsorted populations (P); results from one representative experiment (n=2); error bars indicate SD. (**H**) Clonal capacity of FACS-isolated CD49f^high^ and CD49f^low^ cells from NMU-o-P seeded at 1-100 cells/drop Matrigel (*n*=3). The proportion of Matrigel drops showing outgrowth (bars) and the number of organoids/drop (color code) are shown. (**I**) Monoclonal origin of NMU-o established from a mixture of cells from organoids derived from EGFP-and mTomato-expressing mice. ANOVA and Mann-Whitney tests were applied; * p ≤ 0.05, ** p ≤ 0.01; *** p ≤ 0.001. Scale bar, 500μm.

Three-dimensional (3D) organoids have become a powerful tool to study the molecular and cellular basis of epithelial differentiation, allowing consistent culture and perpetuation (Sato et al., 2009). Organoids are derived from stem/progenitor cells capable of self-renewal and self-organization through cell sorting and lineage commitment in an in vivo-like manner (Kretzschmar and Clevers, 2016). The Clevers laboratory has pioneered the establishment of organoids from a wide variety of epithelia, including mouse small intestine (Sato et al., 2009), liver (Huch et al., 2015), prostate (Karthaus et al., 2014), and pancreas (Boj et al., 2015). Organoids facilitate studying tissue biology, modeling disease, drug screening, and establishing a solid basis for regenerative medicine and gene therapy (Fatehullah et al., 2016). The majority of studies have focused on organoids derived from simple epithelia; work to understand urothelial biology and carcinogenesis lags behind. Here, we harness the ability of ITGA6/CD49f-expressing (CD49^high^) urothelial stem cells to self-perpetuate as organoids, identify the critical pathways involved therein, and use transcriptomics to identify the role of the Notch pathway in urothelial differentiation.

## Results

### Normal mouse urothelial organoids can be established and perpetuated

To establish organoids, we isolated and characterized the cell populations present in digests of urothelial scrapings. Flow cytometry analysis using specific antibodies allowed the purification of EpCAM^+^ epithelial cells from non-epithelial cells expressing CD45^+^, CD31^+^, CD140a^+^, or Ter119^+^ (Figure S1A). EpCAM^+^ cells cultured in Matrigel with complete medium (CM, including EGF, LY2157299, Noggin, WNT3A, RSPO1) (Clevers, 2016; Fatehullah et al., 2016) led to growth of multilayered organoids over one week. Organoids could be consistently passaged and maintained in culture uninterruptedly for >1 year with stable morphology and a tendency to become enriched in cells with basal features (Figure S1B-D). To identify critical growth factors, we performed “Leave-one-out” experiments where we removed each CM component individually or in combination. At day 7, we observed a statistically significant reduction of organoid number upon omission of EGF, WNT3A, RSPO1, or WNT3A+RSPO1 (Figure 1B-C). As reported for other tissues, Noggin was not required to establish organoids but it was essential for long-term perpetuation (not shown) (Yin et al., 2014). Despite their high proliferative potent, organoids did not form tumors upon xenotransplantation (not shown).

### CD49f^high^ expression defines an organoid-forming urothelial cell population with stem cell features

To define the cell-type of origin of the organoids, we isolated urothelial cell subpopulations. Immunofluorescence analysis of normal urothelium showed that CD49f selectively labels basal cells (Figure 1A) (Liu et al., 2012). Purified CD49f^high^ cells were enriched in basal markers whereas CD49f^low^ cells were relatively enriched in luminal markers (Figure 1D). We screened a panel of lectins (not shown) and found that umbrella cells are strongly labeled by Wheat germ agglutinin (WGA), unlike the remaining urothelial cells (Figure 1A). FACS-sorting EpCAM^+^ cells using these markers and cell size – which diminishes towards the luminal cells-showed that CD49f^high^/WGA^+^ cells (basal) have highest organoid-forming capacity. CD49f^low^ cells (intermediate and luminal) were essentially unable to form organoids, regardless of WGA labeling or size (Figure 1E). CD44 has been proposed as a urothelial stem/basal cell marker (Chan et al., 2009; Ho et al., 2012): most CD44^+^ cells were CD49f^+^ and both CD49f^high^/CD44^high^ and CD49f^high^/CD44^low^ cells formed organoids with a similar phenotype (Figure F, S1E, and S1F). However, the number of organoids was significantly higher in the CD49f^high^/CD44^high^ population (P=0.029) (Figure 1G). Low-density seeding and mixing experiments using EGFP- and Tomato-labeled cells showed the monoclonality of the organoids (Figure 1H-I). Altogether, these results indicate that CD49f^high^ labels the organoid-forming urothelial cell population.

### NMU-o recapitulate the urothelial cyto-architecture and display plasticity

To induce differentiation, organoids that had been cultured for 7 days in CM (NMU-o-P) were cultured in differentiation medium (DM) (NMU-o-D) (without EGF, LY2157299, Noggin, WNT3A, and RSPO1) for an additional 7 days: a dramatic morphological change was observed, including a reduced number of cell layers and an increase in lumen formation, indicative of epithelial barrier function (Figure 2A). NMU-o-P expressed high levels of *Ccne1* transcripts and KI67 and resemble basal cells expressing *Cd49f*, *Cd44*, *Tp63*, *Krt14* and *Krt5* and low levels of uroplakins (Figure 2B-C). By contrast, NMU-o-D showed marked downregulation of cell cycle mRNAs and proteins, a modestly decreased expression of basal markers, and high mRNA expression of *Foxa1* and *Ppar*γ, intermediate markers (*Krt8* and *Krt18*) (not shown), and uroplakins (*Upk*2 and *Upk3a*) (Figure 2B). The corresponding proteins displayed the canonical distribution observed in normal urothelium: TP63 and CD49f were found in the outer layer of NMU-o-P while PPARγ and UPK3A were detected in cells lining the lumen of NMU-o-D (Figure 2C). Expression of KRT14 and KRT5 persisted in NMU-o-D, possibly reflecting the half-life of these proteins and the slow differentiation dynamics of urothelial cells in tissues.

**Figure 2.**
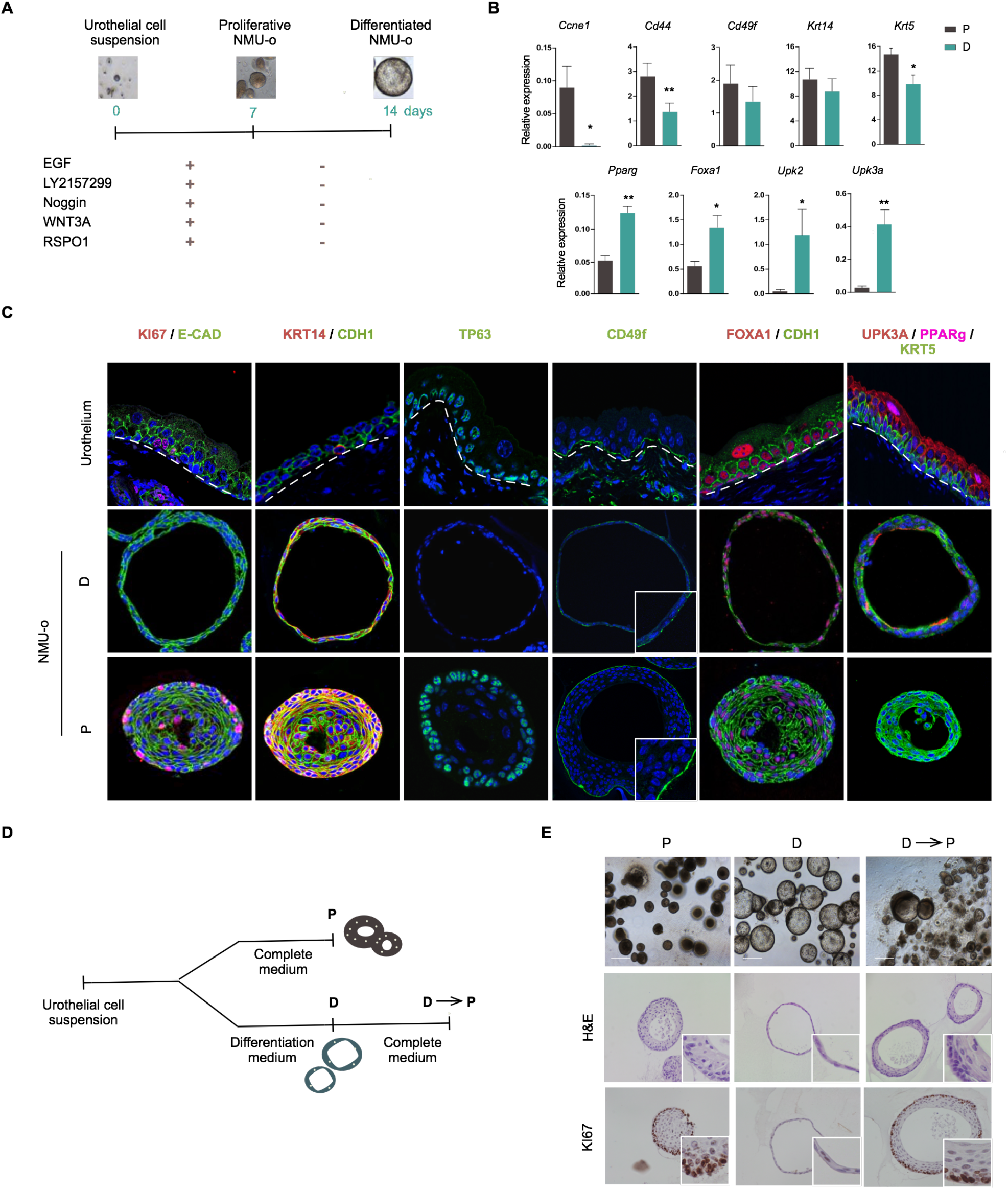
Growth factor-depleted NMU-o recapitulate the urothelial differentiation program and exhibit plasticity. (**A**) Experimental design to induce urothelial differentiation: day 7 NMU-o-P were maintained for 7 additional days in medium lacking CM components. (**B**) RT-qPCR analysis of expression of genes regulated during differentiation. Error bars indicate SD. ANOVA and Mann-Whitney tests were applied; * p ≤ 0.05, ** p ≤ 0.01; Scale bar, 500μm. (**C**) Immunofluorescence analysis of urothelial differentiation markers in proliferative and differentiated NMU-o. DAPI staining is shown in blue. **(D)** Experimental strategy to assess plasticity: day 7 NMU-o-P were maintained for 7 additional days either in complete (P) or differentiation (D) medium. NMU-o-D were then switched to CM (D-P) for 7 additional days. **(E)** Representative phase contrast images of NMU-o cultures, H&E staining, and KI67 expression (n=1).

To determine whether NMU-o-D display terminal differentiation, we cultured them for an additional 7 days either in CM or in DM (Figure 2D-E and S2). NMU-o-D cultured in CM (D→P) acquired a multilayer organization similar to that of NMU-o-P with re-expression of KI67, indicating that they contain cells able to respond to growth cues and undergo dedifferentiation/plasticity (Figure 2E and S2B). Accordingly, 88.9% of cells in NMU-o-D were CD49f^high^ (Figure S2C). NMU-o-D maintained in DM (D→D) showed highest mRNA expression of *Foxa1*, *Pparγ*, and uroplakins (not shown); however, smaller lumina and higher cell death were noted (Figure S2B). These results indicate that NMU-o acquire a cyto-organization and differentiation characteristic of normal urothelium, while retaining progenitor features.

### Combined EGFR inhibition and PPARγ activation drive urothelial organoid differentiation

We used the PPARγ agonist Roziglitazone (Rz) and the EGFR inhibitor Erlotinib to determine if they could further induce urothelial differentiation. NMU-o-P cultured for an additional 7 days in the presence of Rz+Erlotinib acquired larger lumina and showed significantly lower KI67 labeling and higher UPK3A expression, compared to untreated NMU-o-P (Figure 3A-B). *Cd49f* and *Cd44* mRNAs were down-regulated while uroplakin transcripts were up-regulated (Figure 3C). In NMU-o-D, Rz or Erlotinib alone caused reduced expression of *Cd49f* and *Cd44* mRNAs (Figure S3B). When combined, they led to highest *Foxa1* and uroplakin mRNAs expression and to a significant reduction of lumen formation. There were no major changes in KI67 and cleaved-caspase 3 labeling (Figure 3A-C). Treatment of NMU-o-D with the PPARγ inverse agonist T0070907 had minor effects on lumen formation, KI67, and UPK3A expression (Figure 3A-C). KRT20 was not detected in any of the conditions analyzed. These results indicate that PPARγ activation, together with EGFR inhibition, effectively promotes urothelial differentiation and suggest that other pathways contribute to this process.

**Figure 3.**
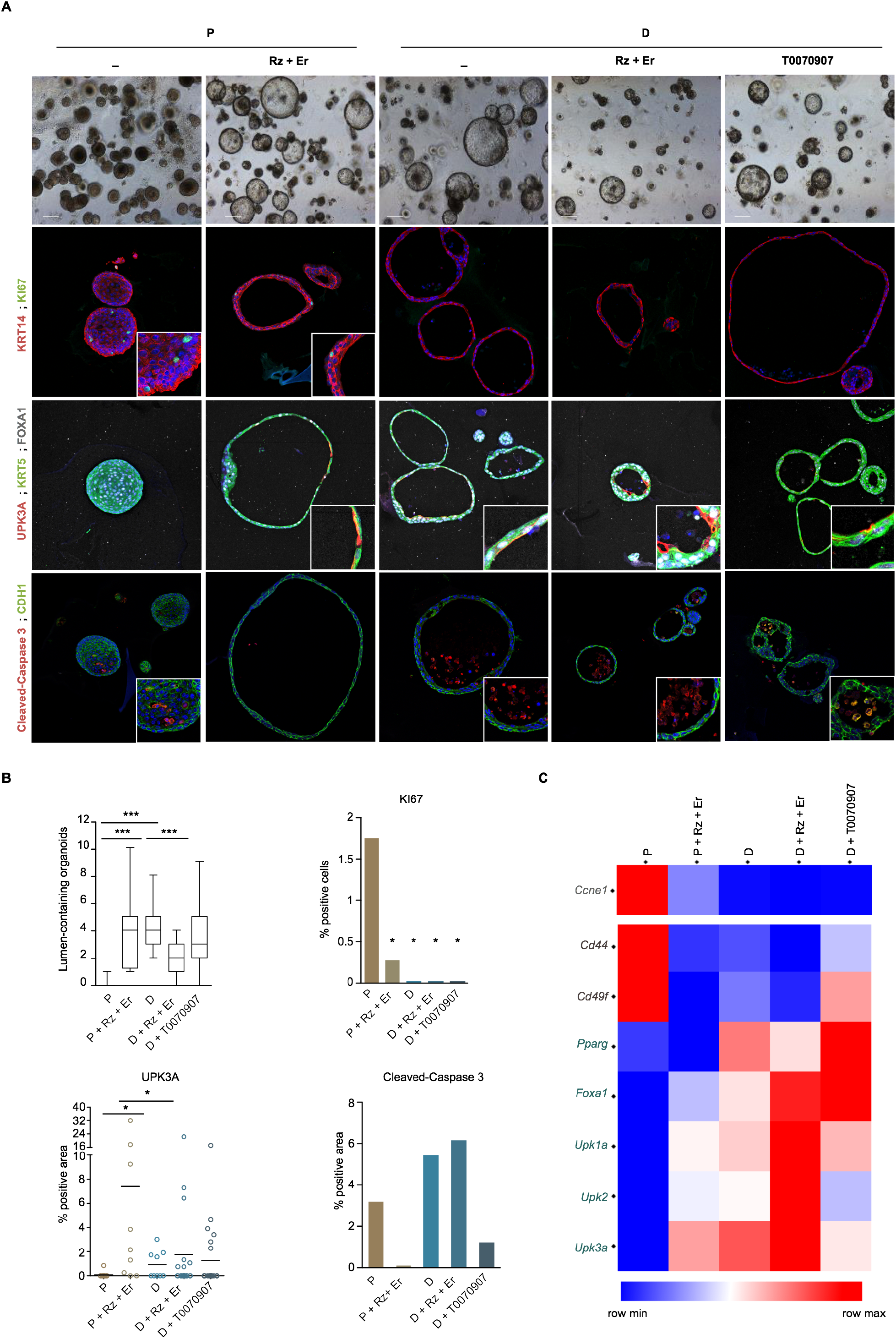
Pharmacological modulation of EGFR and PPARγ activity potentiates NMU-o differentiation. (**A**) Representative phase contrast and immunofluorescence images of proliferative (P) and differentiated (D) NMU-o cultured in the presence of drugs modulating PPARγ and EGFR activity. (**B**) Quantification of lumen formation, KI67, UPK3A, and cleaved caspase 3 in NMU-o cultured as described in panel A. (**C**) Heatmap representing RT-qPCR expression analysis of cell cycle and canonical urothelial differentiation makers in proliferative (P) or differentiated (D) NMU-o treated with Rz+Erlotinib, and with the PPARγ inverse agonist T0070907 (n=2). * p ≤ 0.05, ** p ≤ 0.01; *** p ≤ 0.001. Scale bar, 500μm.

### Transcriptome analysis of differentiated NMU-o reveals novel pathways activated during urothelial differentiation

To interrogate the transcriptomic programs governing NMU-o differentiation we performed RNA-seq of 3 independent paired P and D organoid cultures (Figure S4A). Principal component analysis showed that NMU-o-P clustered closely whereas NMU-o-D showed greater transcriptome divergence (Figure S4B); 4100 genes were differentially expressed between both conditions (FDR<0.05; Table S1). Among the top up-regulated transcripts in NMU-o-P were those involved in cell cycle (i.e. *Cdk1*, *Aurkb*, *Ccnb2*, and *Ki67*), inhibition of apoptosis (*Birc5*), epidermal differentiation (i.e. *Sprr2f*, *Crnn*, *Stfa3*), cytoskeletal regulation (*Kif15*, *Kif4*), and stemness (*Cd34*). By contrast, transcripts significantly up-regulated in NMU-o-D included those involved in urothelial differentiation (i.e. *Upk1a*, *Upk2, Upk3a*), xenobiotic metabolism (i.e. *Cyp2f2*, *Adh7*, *Gstm1*), glycosylation (i.e. *Wbscr17*, *Ugt2b34*, *Galnt14*), and TGF-β signaling (i.e. *Fstl1*, *Ltbp1*, *Fstl4*) (Table S1). Manual curation revealed the down-regulation of canonical basal urothelial markers (i.e. *Krt14*, *Krt5*, *Cd44*, *Cd49f*, and *Tpr63*) and the up-regulation of suprabasal markers (i.e. *Krt19*, *Krt8*, *Foxa1, Ppar*γ *Upk1a*, *Upk1b*, *Upk2*, and *Upk3a*) in NMU-o-D (Figure 4A and S4C); robust regulation of additional keratin species was noted (Figure S4D).

**Figure 4.**
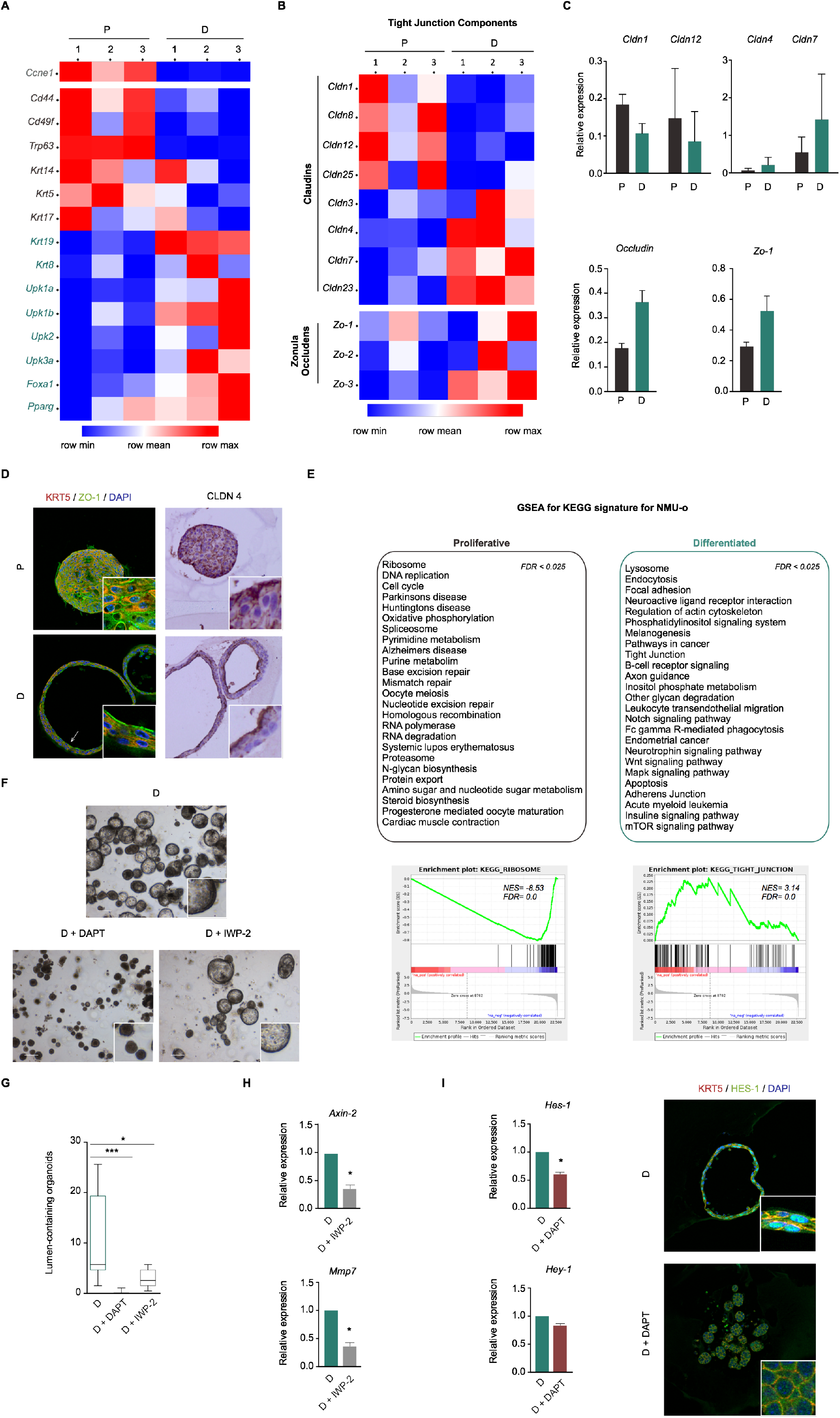
Transcriptome analysis reveals the differentiation capacity of NMU-o, novel pathways activated during differentiation, and the critical role of Notch signaling. **(A)** Heatmap showing the expression (FPKM, RNA-seq) of key urothelial differentiation genes in proliferative and differentiated NMU-o (n=3). **(B)** Heatmap showing the expression of genes related to tight junctions (claudins, occludin, and Zo proteins) (FPKM, RNA-seq) in proliferative and differentiated NMU-o (*n*=3). **(C)** RT-qPCR validation of expression of specific claudin genes in independent organoids (*n*=2); error bars indicate SD. **(D)** Double immunofluorescence analysis of the expression of ZO-1 and KRT5 in proliferative and differentiated NMU-o; immunohistochemical analysis of CLDN4 in the same samples. (**E)** GSEA showing the top 25 pathways (Kyoto Encyclopedia of Genes) significantly enriched in proliferative and differentiated NMU-o. Representative GSEA enrichment plots for the “Ribosome” and “Tight junction” pathways; NES, normalized enrichment scores. **(F-J)** Effect of pharmacological inhibition of Notch (DAPT) and Wnt (IWP-2) on differentiated NMU-o (n=5). Phase contrast (**F**); quantification of lumen formation (**G**); RT-qPCR analysis of Wnt (H) and Notch (I) target genes (**H**); and immunofluorescence for HES-1 (**I**). Error bars indicate SD. * p ≤ 0.05, ** p ≤ 0.01; *** p ≤ 0.001. Scale bar, 500μm.

Epithelial impermeability and endocytic traffic are major features of functional urothelium and both pathways were significantly up-regulated in NMU-o-D. Manual curation revealed dynamic changes in expression of genes coding for proteins involved in cell-cell adhesion. Transcripts coding for tight junction components showed two distinct expression patterns: *Cldn1*, *Cldn8*, *Cldn12*, and *Cldn25* were down-regulated upon differentiation whereas *Cldn3*, *Cldn4*, *Cldn7*, *Cldn23*, *Ocln, Zo-1, Zo-2* and *Zo-3* were up-regulated (Figure 4B), suggesting distinct functions and cellular distribution for the corresponding proteins. Selected mRNA expression changes were confirmed in independent organoid samples and normal urothelium (Figure 4C-D and S4E). The predicted expression pattern of CLDN1, CLDN3, and CLDN4 was validated using the Human Protein Atlas (Figure S4F, https://www.proteinatlas.org/).

Promoter motif analysis of differentially expressed genes using ISMARA revealed significant enrichment in motifs of TF involved in proliferation (E2F, MYC, MYB) and hypoxia regulation (HIF1A) in NMU-o-P. By contrast, NMU-o-D showed enrichment of chromatin regulators (EZH2, IKZF1, CHD1), Hippo pathway (TEAD3, TEAD4), and epithelial-mesenchymal transition regulators (SNAI1, ZEB1, ZEB2) (Figure S4G). Gene set enrichment analysis (GSEA) revealed a highly significant down-regulation of the activity of pathways involved in cell cycle, DNA repair, RNA biology, protein synthesis, and cytokine biology in NMU-o-D; this was accompanied by increased activity of epithelial differentiation/cell-cell adhesion, intracellular traffic (endocytosis, lysosome), and signaling (Notch, phosphatidylinositol, Wnt, MAPK, mTOR) (Figure 4E and Table S2). The porcupine inhibitor IWP-2 reduced Wnt pathway activity and had a modest effect on organoid morphology (Figure 4F-H). Importantly, the γ-secretase inhibitor DAPT significantly reduced expression of Notch target genes and *Upk3a*, and it completely abrogated lumen formation (Figure 4F-G,I, and not shown), supporting the notion that this pathway plays a crucial role in urothelial differentiation.

## Discussion

The identification of normal urothelial stem cells and establishment of conditions for their self-renewal and differentiation is key to an improved understanding of homeostasis and dysregulation in disease. We show that Cd49f labels a subpopulation of basal urothelial cells with stem cell properties and long-term growth potential that largely recapitulates urothelial differentiation, allowing us to uncover new pathways involved therein and providing an important resource for cell biology studies and approaches to regenerative medicine. Alpha2 beta1 and alpha3 beta1 integrins are also markers of skin epidermal stem cells (Jones et al., 1995), underscoring that similar hierarchies exist in stratified epithelia despite fundamental differences in proliferation dynamics. Further analysis of Cd49f^high^ cell heterogeneity should help identify subpopulations with highest self-renewal potential.

There is a remarkable parallelism between the crucial growth pathways involved in urothelial stem cell renewal and those involved in bladder cancer. NMU-o displaying a basal phenotype are critically dependent on EGF for growth and basal/squamous-like bladder tumors show EGFR pathway deregulation, mainly through *EGFR* amplification (Rebouissou et al., 2014). Wnt ligands are required for optimal urothelial organoid growth, consistent with *in vivo* data indicating that stromal Wnt induced by epithelial Shh is required for urothelial recovery from damage impinged by infection or chemical injury (Shin et al., 2011). RNA-seq analysis showed up-regulation of Wnt factor mRNA expression in NMU-o-D and Wnt pathway inhibition reduced the number of lumen-containing organoids, suggesting that epithelial-autonomous Wnt production could contribute to urothelial differentiation. In bladder cancer, genes involved in Wnt signaling are rarely mutated but are frequently deregulated through deletion and/or epigenetic regulation (Stoehr et al., 2004). A role for Wnt has been proposed on the basis of the cooperation of mutant beta-catenin with *Pten* deletion or *HRas* activation in mice (Ahmad et al., 2011a, 2011b).

We show unequivocally that NMU-o undergo differentiation as determined by the expression of urothelial signature markers, as well as by global transcriptomic analysis. Growth factor removal led to the down-regulation of basal signature genes, including epidermal skin transcripts, and the up-regulation of uroplakins and tight junction components. The latter are required for barrier function in the bladder and their up-regulation in NMU-o-D provides evidence of the activation of a physiologically relevant gene expression program. Interestingly, RNA-Seq unveiled differential expression of transcripts coding for members of the Claudin family in NMU-o-P and NMU-o-D, suggesting distinct functions in urothelial cell subpopulations and providing additional differentiation markers. NMU-o-D represent a late – but incomplete – differentiation state as indicated by the lack of KRT20 expression. We speculate that multinucleation, mechanical forces involved in bladder wall distension, and exposure to urine may additionally contribute to regulate KRT20 expression.

PPARγ has been proposed to play a key role in urothelial differentiation. Previous studies using 2D cultures had shown that cells with self-propagating capacity lost the ability to respond to PPARγ agonists (Georgopoulos et al., 2011). By contrast, the organoids we describe here undergo differentiation upon culture with PPARγ activators and EGFR inhibitors in CM. Organoids retained differentiated features in the absence of CM components and upon inhibition of PPARγ with an inverse agonist, suggesting either residual PPARγ activity or the participation of additional signaling pathways. Transcriptomic analysis of NMU-o pointed to a previously unknown role of Notch in normal urothelial differentiation that was confirmed using a gamma-secretase inhibitor. These findings are consistent with genetic evidences indicating that somatic mutations in genes coding for Notch pathway components occur in urothelial tumors (Balbás-Martínez et al., 2013; Rampias et al., 2014; Robertson et al., 2017) and down-regulation of *NOTCH1* and *DLL1* transcript levels in bladder cancer cells (Greife et al., 2014). Notch activation can suppress bladder cancer cell proliferation by direct up-regulation of dual-specificity phosphatases; accordingly, ERK1 and ERK2 phosphorylation was associated with *NOTCH* inactivation and tumor aggressiveness (Rampias et al., 2014). The findings in human samples are supported by genetic mouse models: ubiquitous or urothelium-specific inactivation of Nicastrin, a gamma secretase complex component, led to basal-like tumors and this phenotype was suppressed by inducible overexpression of Notch-IC (Rampias et al., 2014); in addition, urothelial deletion of Presenilin-1/2 or Rbpj accelerated BBN-induced squamous tumors with of epithelial-to-mesenchymal features (Maraver et al., 2015). These results indicate that Notch supports differentiation in normal urothelium and that its inactivation is co-opted during carcinogenesis to promote the development of poorly differentiated tumors.

The identification of a urothelial stem cell population able to form organoids with differentiation capacity, together with single cell analyses, should facilitate the study of the molecular pathophysiology of bladder diseases, including the interaction of epithelial cells with pathogens, and the dissection of the molecular mechanisms through which oncogenes and tumor suppressors contribute to bladder cancer.

## Abbreviations

CM: complete medium
DM: differentiation medium
NMU-o: normal mouse urothelial organoids
Rz: rosiglitazone

## Acknowledgements

We thank the members of the Epithelial Carcinogenesis Group for valuable discussions; N. del Pozo, Y. Gonzalo and R. Serrano for help with animal experimentation; E. Carrillo-de-Santa-Pau and O. Graña for help with bioinformatics analyses; E. Osinaga for help with lectin assays; A. García-España for providing antibodies; D. Chondronasiou, O. Domínguez, J. Gómez-Alonso, N. Malats, L. Martínez, J. L. Martínez-Torrecuadrada, M. Deblas, M. Pérez Martínez, M. Ricote, M. Blasco and M. Serrano, for other valuable contributions; and J. Paramio, P. Martinelli and M. Marqués for critical comments to the manuscript. We thank E. Batlle and his group for many valuable suggestions. This work was supported, in part, by grants from Spanish Ministry of Economy, Industry and Competitiveness (SAF2016-76377-R), CIBERONC (CB16/12/00273) and Fundación Científica de la Asociación Española Contra el Cáncer.

## Author Contributions

Conceptualization, CPS, EL, FXR; Methodology, CPS, EL, LAE, AFB, AB, DM, AM and FXR; Investigation, CPS, EL, LAE; Data Curation, CPS, EL, DM, FXR; Writing – Original Draft, CPS and FXR; Writing – Review & Editing, CPS, EL, AFB, AB, AM, FXR; Funding Acquisition, FXR; Resources, FXR.

---

**Figure S1.**
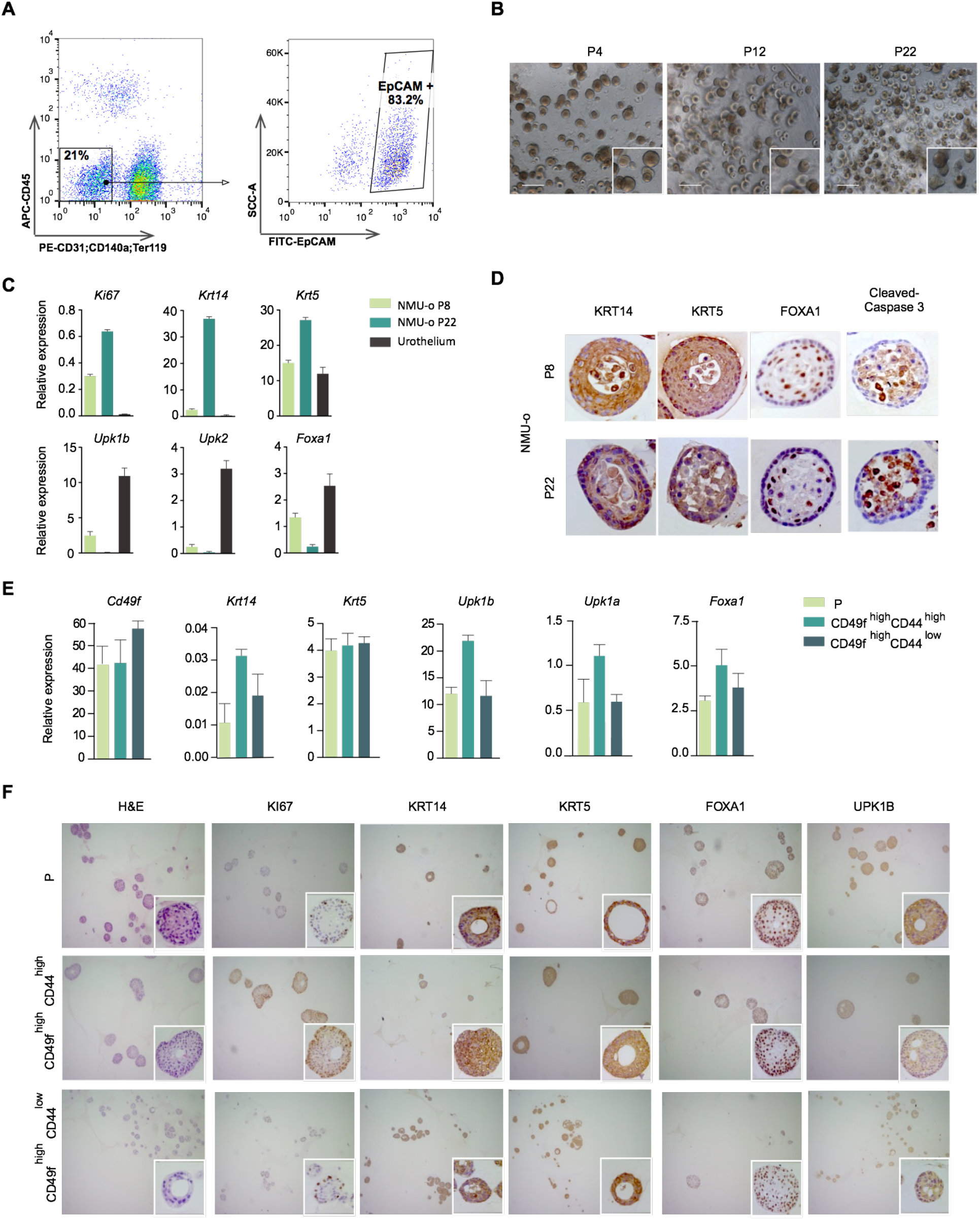

**Figure S2.**
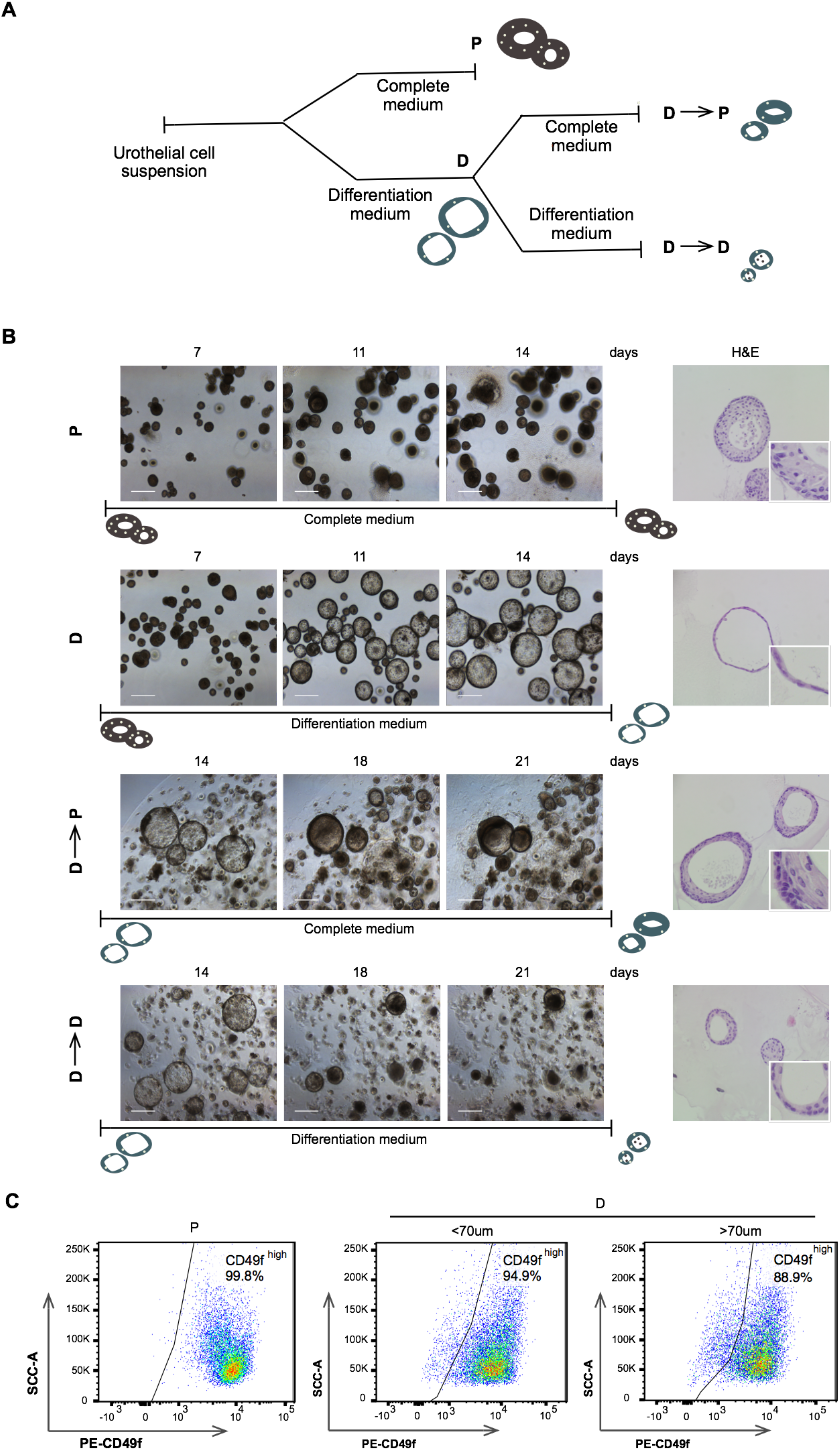

**Figure S3.**
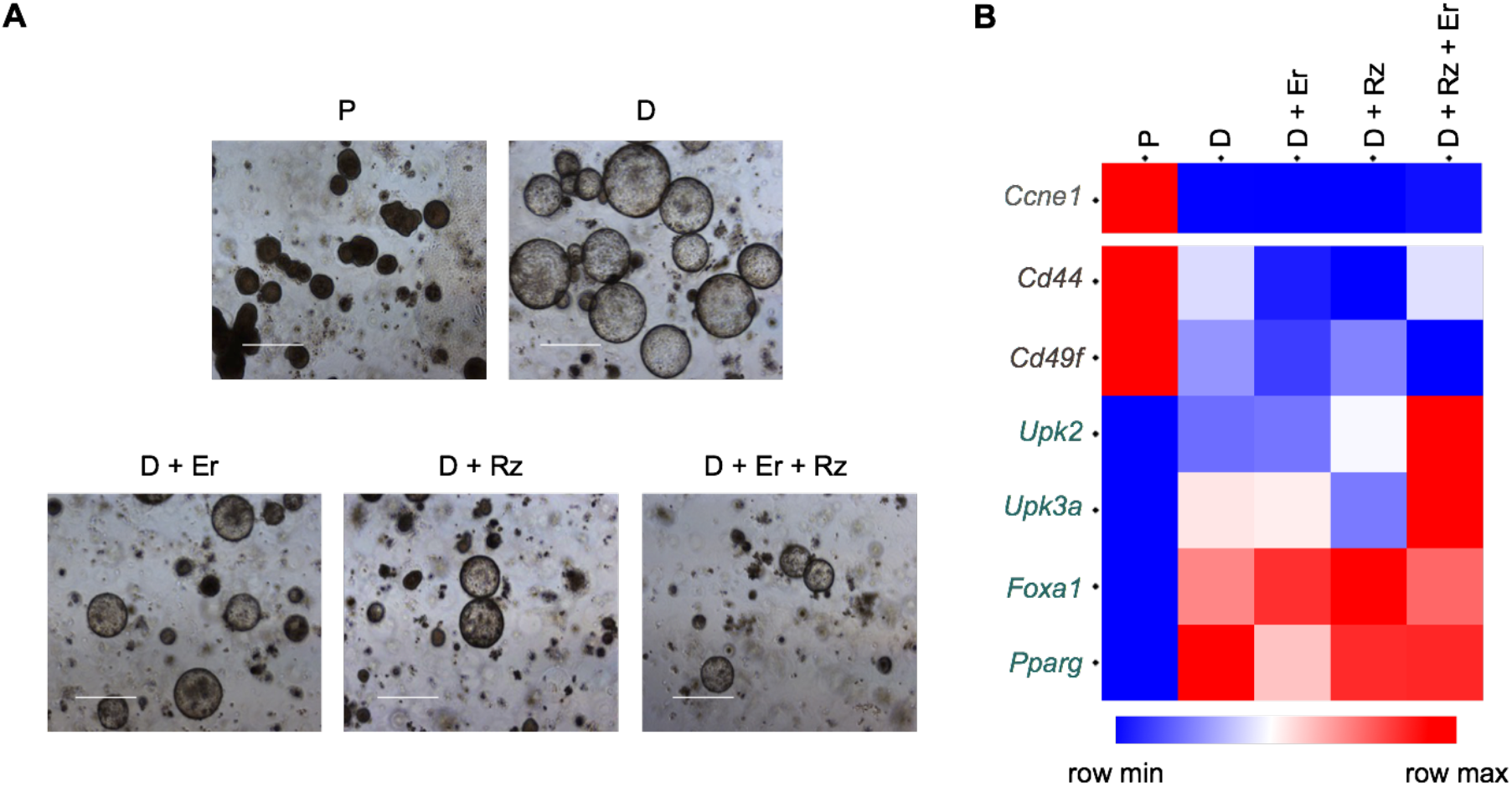

**Figure S4.**
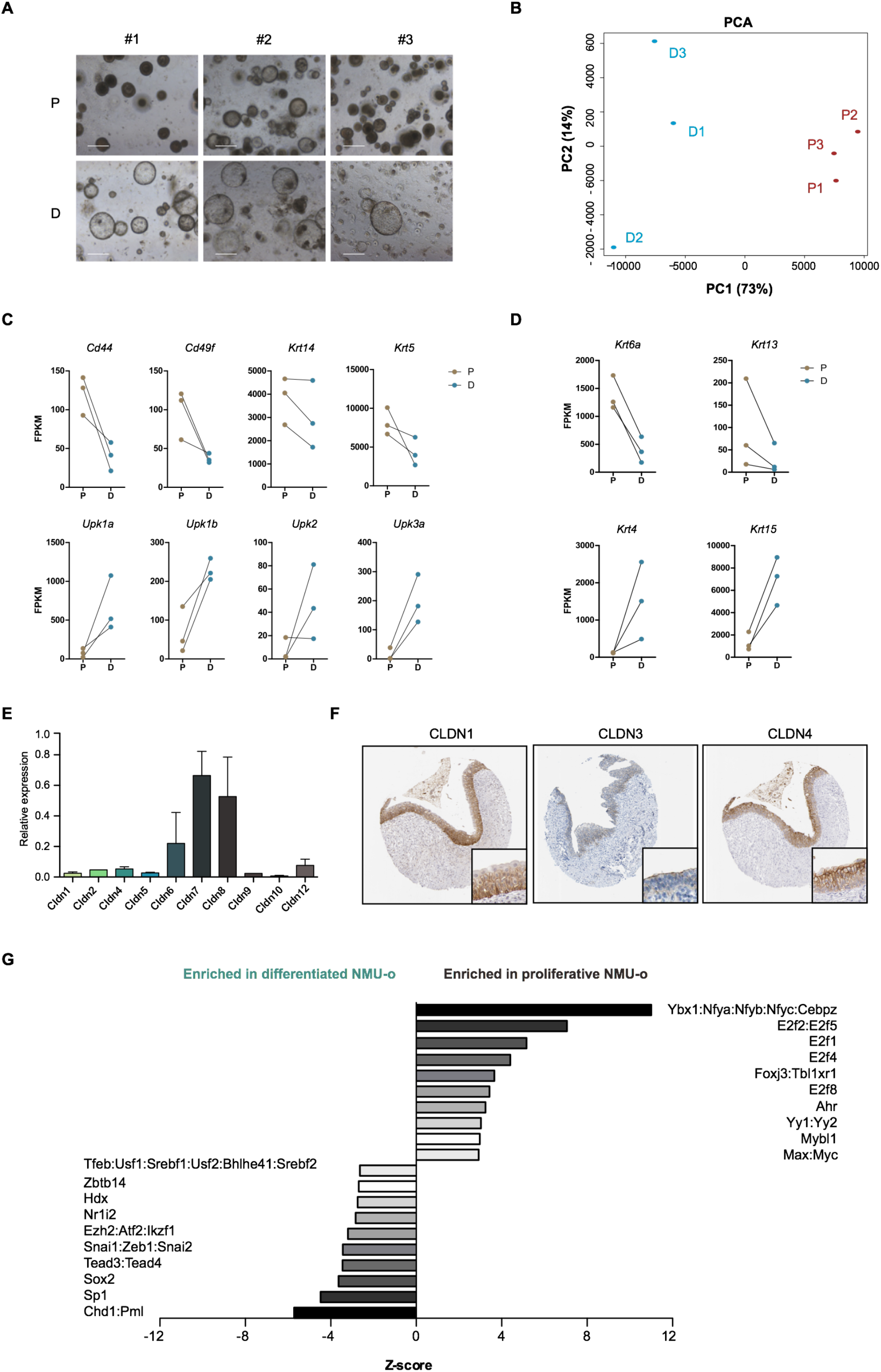

## STAR+METHODS

### KEY RESOURCE TABLE

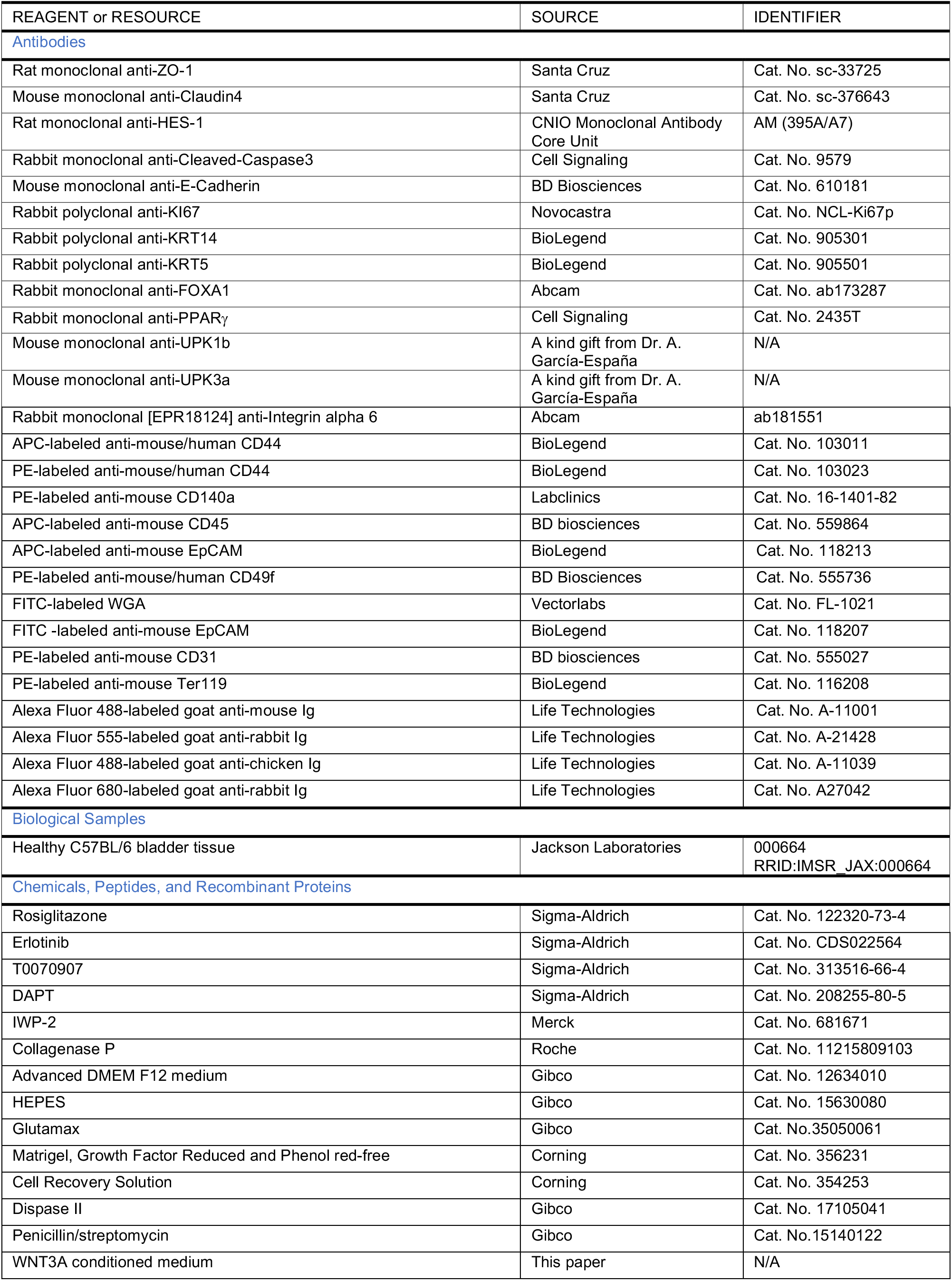
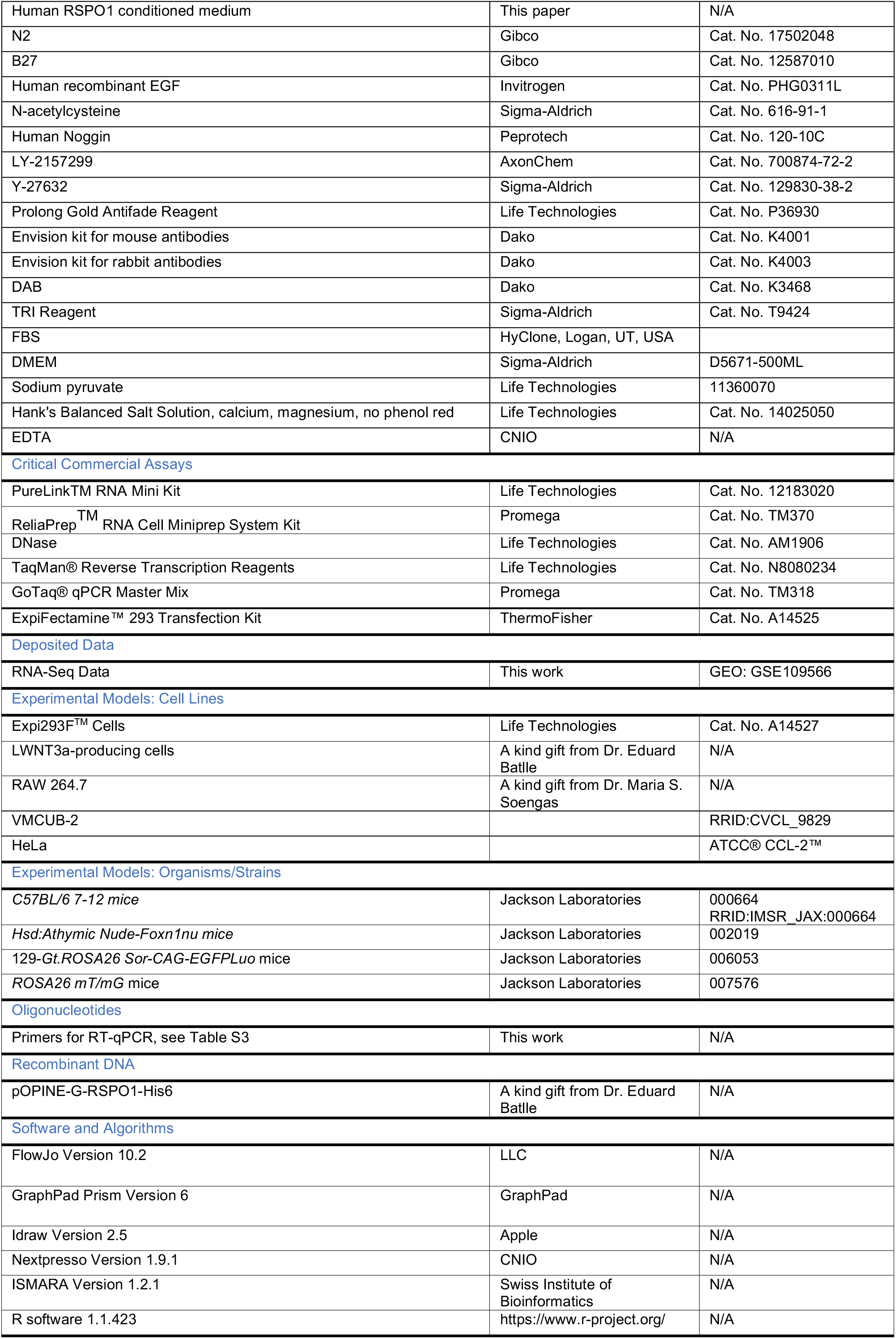
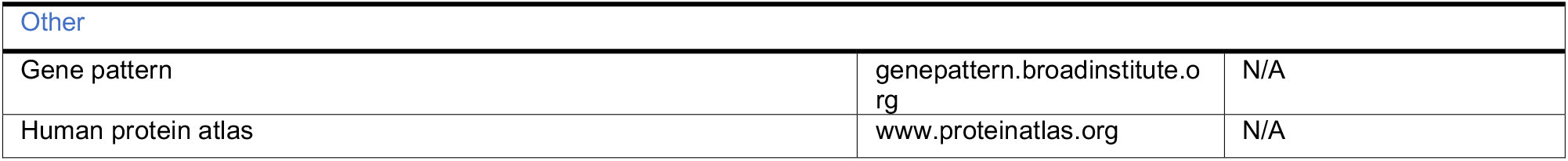

| REAGENT or RESOURCE | SOURCE | IDENTIFIER |
| --- | --- | --- |
| **Antibodies** |  |  |
| Rat monoclonal anti-ZO-1 | Santa Cruz | Cat. No. sc-33725 |
| Mouse monoclonal anti-Claudin4 | Santa Cruz | Cat. No. sc-376643 |
| Rat monoclonal anti-HES-1 | CNIO Monoclonal Antibody Core Unit | AM (395A/A7) |
| Rabbit monoclonal anti-Cleaved-Caspase3 | Cell Signaling | Cat. No. 9579 |
| Mouse monoclonal anti-E-Cadherin | BD Biosciences | Cat. No. 610181 |
| Rabbit polyclonal anti-KI67 | Novocastra | Cat. No. NCL-KI67p |
| Rabbit polyclonal anti-KRT14 | BioLegend | Cat. No. 905301 |
| Rabbit polyclonal anti-KRT5 | BioLegend | Cat. No. 905501 |
| Rabbit monoclonal anti-FOXA1 | Abcam | Cat. No. ab173287 |
| Rabbit monoclonal anti-PPARγ | Cell Signaling | Cat. No. 2435T |
| Mouse monoclonal anti-UPK1b | A kind gift from Dr. A. García-España | N/A |
| Mouse monoclonal anti-UPK3a | A kind gift from Dr. A. García-España | N/A |
| Rabbit monoclonal [EPR18124] anti-Integrin alpha 6 | Abcam | ab181551 |
| APC-labeled anti-mouse/human CD44 | BioLegend | Cat. No. 103011 |
| PE-labeled anti-mouse/human CD44 | BioLegend | Cat. No. 103023 |
| PE-labeled anti-mouse CD140a | Labclinics | Cat. No. 16-1401-82 |
| APC-labeled anti-mouse CD45 | BD biosciences | Cat. No. 559864 |
| APC-labeled anti-mouse EpCAM | BioLegend | Cat. No. 118213 |
| PE-labeled anti-mouse/human CD49f | BD Biosciences | Cat. No. 555736 |
| FITC-labeled WGA | Vectorlabs | Cat. No. FL-1021 |
| FITC -labeled anti-mouse EpCAM | BioLegend | Cat. No. 118207 |
| PE-labeled anti-mouse CD31 | BD biosciences | Cat. No. 555027 |
| PE-labeled anti-mouse Ter119 | BioLegend | Cat. No. 116208 |
| Alexa Fluor 488-labeled goat anti-mouse Ig | Life Technologies | Cat. No. A-11001 |
| Alexa Fluor 555-labeled goat anti-rabbit Ig | Life Technologies | Cat. No. A-21428 |
| Alexa Fluor 488-labeled goat anti-chicken Ig | Life Technologies | Cat. No. A-11039 |
| Alexa Fluor 680-labeled goat anti-rabbit Ig | Life Technologies | Cat. No. A27042 |
| **Biological Samples** |  |  |
| Healthy C57BL/6 bladder tissue | Jackson Laboratories | 000664<br>RRID:IMSR_JAX:000664 |
| **Chemicals, Peptides, and Recombinant Proteins** |  |  |
| Rosiglitazone | Sigma-Aldrich | Cat. No. 122320-73-4 |
| Erlotinib | Sigma-Aldrich | Cat. No. CDS022564 |
| T0070907 | Sigma-Aldrich | Cat. No. 313516-66-4 |
| DAPT | Sigma-Aldrich | Cat. No. 208255-80-5 |
| IWP-2 | Merck | Cat. No. 681671 |
| Collagenase P | Roche | Cat. No. 11215809103 |
| Advanced DMEM F12 medium | Gibco | Cat. No. 12634010 |
| HEPES | Gibco | Cat. No. 15630080 |
| Glutamax | Gibco | Cat. No.35050061 |
| Matrigel, Growth Factor Reduced and Phenol red-free | Corning | Cat. No. 356231 |
| Cell Recovery Solution | Corning | Cat. No. 354253 |
| Dispase II | Gibco | Cat. No. 17105041 |
| Penicillin/streptomycin | Gibco | Cat. No.15140122 |
| WNT3A conditioned medium | This paper | N/A |
| Human RSPO1 conditioned medium | This paper | N/A |
| N2 | Gibco | Cat. No. 17502048 |
| B27 | Gibco | Cat. No. 12587010 |
| Human recombinant EGF | Invitrogen | Cat. No. PHG0311L |
| N-acetylcysteine | Sigma-Aldrich | Cat. No. 616-91-1 |
| Human Noggin | Peprtech | Cat. No. 120-10C |
| LY-2157299 | AxonChem | Cat. No. 700874-72-2 |
| Y-27632 | Sigma-Aldrich | Cat. No. 129830-38-2 |
| Prolong Gold Antifade Reagent | Life Technologies | Cat. No. P36930 |
| Envision kit for mouse antibodies | Dako | Cat. No. K4001 |
| Envision kit for rabbit antibodies | Dako | Cat. No. K4003 |
| DAB | Dako | Cat. No. K3468 |
| TRI Reagent | Sigma-Aldrich | Cat. No. T9424 |
| FBS | HyClone, Logan, UT, USA |  |
| DMEM | Sigma-Aldrich | D5671-500ML |
| Sodium pyruvate | Life Technologies | 11360070 |
| Hank's Balanced Salt Solution, calcium, magnesium, no phenol red | Life Technologies | Cat. No. 14025050 |
| EDTA | CNIO | N/A |
| **Critical Commercial Assays** |  |  |
| PureLink<sup>TM</sup> RNA Mini Kit | Life Technologies | Cat. No. 12183020 |
| ReliaPrep<sup>TM</sup> RNA Cell Miniprep System Kit | Promega | Cat. No. TM370 |
| DNase | Life Technologies | Cat. No. AM1906 |
| TaqMan<sup>®</sup> Reverse Transcription Reagents | Life Technologies | Cat. No. N8080234 |
| GoTaq<sup>®</sup> qPCR Master Mix | Promega | Cat. No. TM318 |
| ExpiFectamine<sup>TM</sup> 293 Transfection Kit | ThermoFisher | Cat. No. A14525 |
| **Deposited Data** |  |  |
| RNA-Seq Data | This work | GEO: GSE109566 |
| **Experimental Models: Cell Lines** |  |  |
| Expi293F<sup>TM</sup> Cells | Life Technologies | Cat. No. A14527 |
| LWNT3a-producing cells | A kind gift from Dr. Eduard Batlle | N/A |
| RAW 264.7 | A kind gift from Dr. Maria S. Soengas | N/A |
| VMCUB-2 |  | RRID:CVCL_9829 |
| HeLa |  | ATCC<sup>®</sup> CCL-2<sup>TM</sup> |
| **Experimental Models: Organisms/Strains** |  |  |
| C57BL/6 7-12 mice | Jackson Laboratories | 000664<br>RRID:IMSR_JAX:000664 |
| Hsd:Athymic Nude-Foxn1nu mice | Jackson Laboratories | 002019 |
| 129-Gt.ROSA26 Sor-CAG-EGFP<sup>Luo</sup> mice | Jackson Laboratories | 006053 |
| ROSA26 mT/mG mice | Jackson Laboratories | 007576 |
| **Oligonucleotides** |  |  |
| Primers for RT-qPCR, see Table S3 | This work | N/A |
| **Recombinant DNA** |  |  |
| pOPINE-G-RSPO1-His6 | A kind gift from Dr. Eduard Batlle | N/A |
| **Software and Algorithms** |  |  |
| FlowJo Version 10.2 | LLC | N/A |
| GraphPad Prism Version 6 | GraphPad | N/A |
| Idraw Version 2.5 | Apple | N/A |
| Nextpresso Version 1.9.1 | CNIO | N/A |
| ISMARA Version 1.2.1 | Swiss Institute of Bioinformatics | N/A |
| R software 1.1.423 | https://www.r-project.org/ | N/A |
| Other |  |  |
| Gene pattern | genepattern.broadinstitute.org | N/A |
| Human protein atlas | www.proteinatlas.org | N/A |

## CONTACT FOR REAGENT AND RESOURCE SHARING

Additional information and requests for resources and reagents should be directed to and will be fulfilled by the Lead Contact, Francisco X. Real.

## EXPERIMENTAL MODEL AND SUBJECT DETAILS

### Mice

C57BL/6, Hsd:Athymic Nude-*Foxn1nu*,129-Gt.*ROSA26 Sor-CAG-EGFPLuo* and *ROSA26 mT/mG* mice (Jackson Laboratories, Bar Harbour, ME) were housed in a specific pathogen-free environment according to institutional guidelines. *129-Gt.ROSA26 Sor-CAG-EGFPLuo* mice were kindly provided by Maria A. Blasco (CNIO). Mice were sacrificed by CO_2_ inhalation. Urothelial bladder suspensions were produced from 7-12 week-old mice both for the establishment of organoids and for fluorescence-activated cell sorting (FACS) analyses. For *in vivo* xenograft transplantation, Hsd:Athymic Nude-*Foxn1nu* mice were injected with 1 million cells/100μL Matrigel per injection. All experiments were performed in accordance with the guidelines for Ethical Conduct in the Care and Use of Animals as stated in The International Guiding Principles for Biomedical Research involving Animals, developed by the Council for International Organizations of Medical Sciences (CIOMS), and were approved by the institutional Ethics and Animal Welfare Committee (Instituto de Salud Carlos III, Madrid, Spain).

### Cell lines

VMCUB-2 cells cultured in basic DMEM were used as a positive control for the xenograft experiments. L-Wnt3a cells transfected with the Wnt3A cDNA were used to produce Wnt3A conditioned medium (Jung et al., 2011). RSPO-1 was supplemented as the conditioned medium of Expi293 cells transfected with a pOPINE-G-RSPO1-His6 plasmid (a kind gift of Dr. E. Batlle, IRB Barcelona) using the ExpiFectamine™ 293 Transfection Kit. The activity of both conditioned media was confirmed by testing their effect on *Axin2* mRNA expression by HeLa cells using RT-qPCR. Murine macrophage RAW 264.7 cells were maintained in Dulbecco’s Modified Eagle’s medium (DMEM), supplemented with 10% FBS, 1% sodium pyruvate and 1% penicillin/streptomycin (basic DMEM). For selected experiments, cells were seeded at 70% confluence in basic DMEM supplemented with oxLDL (15μg/mL) alone or with either Roziglitazone (10μM) or T0070907 (2μM). After 2h, cells were cultured with basic DMEM supplemented with oxLDL (15μg/mL). After 48h, RNA was extracted. All cells were allowed to settle in a humidified incubator at 37 ºC, 5% CO_2_. All cell lines used were Mycoplasma-free.

### Establishment of normal mouse organoids

Mice were sacrificed and the bladder was accessed and turned inside-out leaving the urothelial surface exposed. The urothelium was enzymatically digested with collagenase P (0.5μg/mL) in Hank’s Balanced Salt Solution (HBSS) in a thermoblock with gentle shaking at 37 ºC for 20 min. Collagenase P was inactivated with 2mM EDTA and 50% FBS. The cell suspension was collected and the remaining urothelium was scraped. After filtering through a 70μm strainer (Falcon) and centrifugation at 1200rpm for 5min at 4 ºC, the pellet was washed 2x (Advanced DMEM F12 medium

+ 1x HEPES + 1x Glutamax-washing medium) and cells were embedded in Matrigel (growth factor-reduced and phenol red-free): Matrigel-cell suspensions (20 μL drops) were plated onto 6-well plates, allowed to settle in a humidified incubator at 37 ºC/5% CO_2_, and overlaid with 2mL complete medium (see below), changed every 2-3 days. Cultures were usually expanded at a 1:4-1:6 ratio every 7-10 days. After Matrigel removal with Corning® Cell Recovery Solution on ice, the cell suspension was washed with PBS, then with washing medium, and centrifuged at 1200rpm for 5min at 4 ºC. Then, NMU-o were chemically digested with Dispase II solution (10mg/mL) (Gibco) for 15-20min in a rotating wheel at room temperature. The reaction was neutralized with 2mM EDTA and single cells were obtained by mechanical disruption with a syringe with a 21G needle for at least strokes. After 2 washes with washing medium, the pellet was embedded in fresh ice-cold Matrigel, plated on 6-well plates, and covered with medium as described. Unless otherwise specified, NMU-o were maintained in culture with complete medium [Advanced DMEM/F12, 1x penicillin/streptomycin, 1x HEPES, 1x GlutaMAX, 50% WNT3A conditioned medium, 5% human RSPO1 conditioned medium, 1x N2 (Gibco), 1x B27 (Gibco), 50ng/mL human recombinant EGF (Invitrogen), 1mM N-acetylcysteine (Sigma-Aldrich), 50μg/mL human Noggin (Peprotech) and 1μM LY-2157299 (AxonChem)]. The ROCK inhibitor Y-27632 (10μM) (Sigma-Aldrich) was added during the first 3 days of culture. For differentiation experiments, NMU-o were cultured for the first 7 days in complete medium, reseeded (without disaggregation) in fresh Matrigel, and cultured either with complete medium or with differentiation medium (lacking WNTA3A and RSPO-1 conditioned medium, EGF, LY-2157299 and Noggin) for the following 7 days. PPARγ pathway modulators were added to the medium (complete or differentiated) as indicated in the text, at the following concentrations: Rosiglitazone (1μM), T0070907 (10μM), Erlotinib (0.5μM). The γ-secretase inhibitor DAPT (5μM) and IWP-2 (5μM) were added on day 7 to NMU-o-P and cultures were allowed to grow under differentiation medium until day 14. Microphotographs were taken using an inverted microscope (Olympus CK-30). In order to cryopreserve NMU-o, 7 day cultures were washed once in PBS, Corning® Cell Recovery Solution was added, and cells were collected and placed on ice for 30min. After washing once with PBS, then with washing medium, and centrifugation at 1200 rpm 4 ºC for 5 min, organoids were resuspended in freezing media (10% DMSO in Advanced DMEM/F12 supplemented with 10μM Y-27632) at a density equivalent to 3 confluent drops/500μL. Cryovials were stored at −80 ºC. For thawing, vials were placed in a 37 ºC water bath and the contents washed twice with Advanced DMEM/F12 before reseeding in Matrigel at the required density.

## METHODS DETAILS

### Flow cytometry analysis of isolated urothelial cells and NMU-o

Urothelial cell suspensions obtained as previously described were incubated with blocking buffer (1% BSA/3mM EDTA in PBS) for 15 min at room temperature. After washing twice with PBS, cells were incubated with FITC-labeled anti-EpCAM, APC-labeled anti-EpCAM, FITC-labeled WGA, PE-labeled anti-CD49f, APC-labeled anti-CD45, PE-labeled anti-CD31, PE-labeled anti-CD14a, PE-labeled anti-Ter119 and/or APC-labeled anti-CD44 antibodies in FACS buffer (0.1%BSA/3mM EDTA in PBS) for 30 min at 4 ºC. After washing twice with PBS, cells were resuspended in FACS buffer and stained with DAPI (Sigma-Aldrich). In all the experiments a control sample lacking primary antibody and a Fluorescence Minus One (FMO) control were used. In the experiments from isolated urothelial cells, EpCAM+ single cells were sorted by FACS and dead cells were excluded from subsequent analyses. In the experiments with NMU-o, samples were disaggregated as previously mentioned and single cell suspensions were incubated with PE-labeled anti-CD49f antibodies. In the case of D<70μm and D>70μm, NMU-o-P at day 7 were filtered using a 70μm filter and the fallthrough and the retained organoids were reseeded in Matrigel and cultured in differentiation medium. All samples were analysed using a FACS Influx or AriaII (BD Biosciences) flow cytometer and at least 10,000 events were acquired. Analyses were performed using FlowJo flow cytometer analysis software.

### NMU-o clonality experiments

FACS-sorted cells were embedded in 5μL of Matrigel in a 96-well format at 1, 10 or 100 cells/well for the clonal growth experiments. For the monoclonality experiment, organoids from 129-Gt.*ROSA26 Sor-CAG-EGFPLuo* and from *ROSA26 mT/mG* urothelia were separately established; after dissociation to single cell suspensions they were mixed at a 1:1 ratio and allowed to grow as organoids in 20μL of Matrigel/drop in 48-well plates at 1000 cells/well. Organoids of the corresponding fluorescent colour (EGFP, Tomato, and chimeras) were counted.

### NMU-o plasticity

Urothelial cell suspensions were seeded in complete medium for 7 days. Afterwards, NMU-o were resuspended in Matrigel and cultured for the following 7 days either in complete medium or in differentiation medium. On day 14, the medium of NMU-o-D was changed to either complete or differentiation medium and cultures were allowed to grow until day 21.

### NMU-o quantification

For the “leave-one-out experiments”, images were acquired with X4 resolution with CCD-microscope using a bright-field filter. Three pictures in the Z axis were taken in order to collect the majority of the organoids. Then, a Z-stack was done using ImageJ software. For immunofluorescence and growth assays, quantification was performed with tailored routines programmed in Definiens XD v2.5 software. Organoid growth and time-lapse video of were performed on a DMI6000B bright field microscope from Leica Microsystems by using an HC PL APO 10x 0.4N.A dry objective. Cells were maintained in a temperature-controlled (37 ºC), humidified environment in the presence of 5% CO_2_ during imaging.

### Immunofluorescence (IF) and immunohistochemistry (IHC) staining

Matrigel drops containing organoids were collected 7 days after seeding, fixed in 10% formalin, and embedded in paraffin. Sections (3 μm) were obtained for IF and IHC analyses. After deparaffinization and rehydration, antigen retrieval was performed by boiling in citrate buffer pH 6 for 10 min. For IF, sections were incubated with 3% BSA/0.1% Triton in PBS for 45 min at room temperature and with primary antibody overnight at 4 ºC. After washing with 0.1% Triton/PBS, the appropriate fluorochrome-conjugated secondary antibodies were added for 45 min, sections were washed, and nuclei were counterstained with DAPI (Sigma). After washing with PBS, sections were mounted with Prolong Gold Antifade Reagent (Life Technologies). Images were acquired using a confocal microscope (Leica, SP5) using a x40 immersion oil lens with a zoom between 1 and 2.5. For IHC, endogenous peroxidase was blocked with 3% H_2_O_2_ in PBS for 30 min, sections were incubated with 3% BSA/0.1% Triton in PBS for 45 min at room temperature, and with primary antibody overnight at 4 ºC. After washing with 0.1% Triton/PBS, sections were incubated with the appropriate secondary antibodies (Envision kit for mouse or rabbit Ig, Dako) for 45 min, washed, and reactions were developed with DAB (Dako). Sections were lightly counterstained with haematoxylin, dehydrated, and mounted. The following reagents were used: FITC-labeled WGA, PE-labeled anti-CD44, PE-labeled anti-CD49f, anti-claudin4, anti-Zo-1, anti-PPARγ, anti-KI67, anti-FOXA1, anti-E-Cadherin, anti-Cleaved-caspase3, anti-KRT5, anti-KRT14, anti-TP53, anti-UPK3a, anti-UPK1b, anti-ITGA6/CD49f, and anti-HES1.

### Real-Time quantitative PCR

Total RNA was extracted from organoids using the TRI Reagent® (Sigma-Aldrich) followed by the PureLinkTM RNA Mini Kit (Life Technologies), according to manufacturer’s instructions. For 2D culture, RNA was extracted using ReliaPrep^TM^ RNA Cell Miniprep System Kit (Promega). Samples were treated with DNase before reverse transcription (Life Technologies). cDNA was generated from 200-1000ng of RNA using random hexamers and reverse transcriptase using the TaqMan® Reverse Transcription Reagents (Life Technologies). Reaction mixes lacking RT were used to ensure the absence of genomic DNA contamination. RT-PCR amplification and analyses were conducted using the 7900HT Real-Time PCR System (Applied Biosystems, Life Technologies) using GoTaq® qPCR Master Mix (Promega). Gene-specific expression was normalized to *Hprt* (organoid cultures), *β-actin* (RAW 264.7 cells) and *GAPDH* (HeLa cells) using the ΔΔCt method. For RT–qPCR, primer pairs were designed to achieve inter-exon products of 200–250 bp. Primer sequences are provided in Table S1.

### RNA-seq analysis

RNA quality was assayed by laboratory chip technology on an Agilent 2100 Bioanalyzer. PolyA+ RNA was isolated from total RNA (1μg, RIN>9), randomly fragmented, converted to double stranded cDNA, and processed through subsequent enzymatic treatments of end-repair, dA-tailing and ligation to adapters according to Illumina’s TruSeq RNA Sample Preparation v.2 Protocol. The adaptor-ligated library was completed by limited-cycle PCR with Illumina PE primers (8 cycles). The resulting purified cDNA library was applied to an Illumina flow cell for cluster generation (TruSeq cluster generation kit v.5) and sequenced on a Genome Analyzer IIx with SBS TruSeq v.5 reagents, according to the manufacturer’s protocols. The sample sequencing was paired-end. Three paired NMU-o-P and three NMU-o-D samples were analysed. The Nextpresso analysis pipeline (Bioinformatics Unit, CNIO, Madrid) was used to process the data with the version MGSCv376/mm8 of the mouse genome (Graña et al., 2017).

### Principal component analysis

The Pearson correlation was calculated from the expression values (expressed as fragments per kilobase of transcript per million mapped reads) of each gene for each sample by using the ‘cor’ command in R (https://www.r-project.org/). Principal component analysis was performed using the ‘prcomp’ command in R, from the correlation value of each sample.

### Gene Set Enrichment Analysis

The list of genes was ranked by the ‘t-stat’ statistical value from the cuffdiff output file. The list of pre-ranked genes was then analysed with GSEA for Gene Ontology (GO) database. Significantly enriched GO terms were identified using a false discovery rate q value of less than 0.25. The analyses were carried out as defined in http://www.broadinstitute.org/gsea/doc/GSEAUserGuideFrame.html?Interpreting_GSEA.

## ADDITIONAL RESOURCES

Claudin expression data was extracted from the Human Protein Atlas (www.proteinatlas.org) in order to validate claudin expression in human bladder tissue. GenePattern was used to compute all heatmaps here presented. IDraw was used to generate the panels and figures.

## QUANTIFICATION AND STATISTICAL ANALYSIS

All quantitative data are presented as mean±s.e.m. from ≥2 experiments or samples per data point (*n* is mentioned in each figure legend). Non-parametric Mann–Whitney U test (two-tailed) was used to assess significance levels and ANOVA was used to compare more than 2 groups. Statistical analysis was performed using GraphPad Prism software. For further statistical details, refer to each figure legend. For *in vitro* experiments, sample size required was not determined a priori. The experiments were not randomized.

## DATA AND SOFTWARE AVAILABILITY

The RNA-seq gene expression data generated in this study have been deposited in GEO with accession number GSE109566. Excel files with significantly differentially regulated genes and pathways are available upon request to the authors.

